# Rescoring ensembles of protein-protein docking poses using consensus approaches

**DOI:** 10.1101/2020.04.24.059469

**Authors:** Guillaume Launay, Masahito Ohue, Julia Prieto Santero, Yuri Matsuzaki, Cécile Hilpert, Nobuyuki Uchikoga, Takanori Hayashi, Juliette Martin

## Abstract

Scoring is a challenging step in protein-protein docking, where typically thousands of solutions are generated. Successful scoring is more often based on physicochemical evaluation of the generated interfaces and/or statistical potentials that reproduce known interface properties. Another route is offered by consensus-based rescoring, where the set of solutions is used to build statistics in order to identify recurrent solutions. We explore several ways to perform consensus-based rescoring on the ZDOCK decoy set for Benchmark 4. We show that the information of the interface size is critical for successful rescoring. We combine consensus-based rescoring with the ZDOCK native scoring function and show that this improves the initial results.

## INTRODUCTION

Protein-protein docking aims at predicting the structure of a complex starting from the structures of isolated components ^1, 2^. The CAPRI community-wide initiative allows a blind assessment of the participant methods on common data sets and evaluation criteria, offering an updated view of progress in the field since 2001 ^3–5^. Protein-protein docking methods typically generate thousands of potential solutions for a particular complex. Scoring the models to discriminate near-native solutions is a known bottleneck of docking methods ^6–8^. Most scoring functions are physics-based, attempting to capture the determinants underlying the stability of protein-protein complexes, e.g., shape complementary, electrostatics and desolvation potential ^9–15^. Knowledge-based functions, on the other hand, aim at taking advantage of the information from available structures, *via* pair potentials ^16, 18^, or multibody potentials ^19^. Docking methods often use scoring functions that combine physical terms with knowledge-based terms ^20–23^. More recently, evolutionary information has been successfully used for scoring ^24, 25^.

Another approach consists in relying on the recurrences observed in the set of solutions, i.e., consensus-based scoring. Consensus-based scoring functions seek to identify solutions with features that are the most frequent in the solution set, independently of any physics-based or evolutionary evaluation. The CONSRANK scoring function, proposed by Oliva et al ^26–29^ has shown very good results, based on the conservation of interface contacts.

In this study, we compare several consensus-based scoring functions on large sets of docking poses generated by ZDOCK, including CONSRANK. We then propose a way to combine consensus-based rescoring with the native scoring function of ZDOCK, showing that it is possible to improve initial results.

## METHODS

### Docking decoy set

The ZDOCK3.0.2 decoy set ^30^ (6 degree sampling, fixed receptor format) for Benchmark4 ^31^ was retrieved from https://zlab.umassmed.edu/zdock/decoys.shtml. This data set encompasses 176 protein-protein complexes, with 54,000 docking poses for each complex. For each pose, the interface Cα RMSD, with respect to the bound structure, is given. A near-native docking hit is defined as a prediction with interface Cα RMSD < 2.5 Å.

### Consensus-based rescoring schemes

Following the CONSRANK method ^26, 27^, docking poses are rescored using the frequencies of interface contacts in the set of docking poses. Interface contacts are defined using a distance cut-off of 5 Å between the heavy atoms of receptor and ligand proteins.

For each contact *C_ij_* between residue *i* from receptor and residue *j* from ligand the relative frequency in the decoy set is defined by:

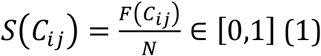

where *F*(*C_ij_*) denotes the frequency of *C_ij_* at the protein-protein interface in the set of *N* decoys. These relative frequencies are averaged to compute the CONSRANK score of each pose P:

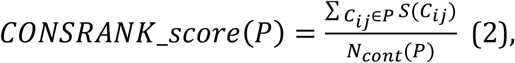

where *N_cont_*(*P*) denotes the number of interface contacts in docking pose *P*.

### Variations of CONSRANK scores

In addition to CONSRANK, we implemented alternative consensus-based scoring functions.

First, we considered an un-normalized version of CONSRANK scores, where the sum of contact contributions is not averaged by the interface size:

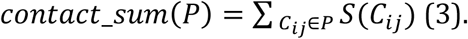

Then, we implemented two other variations, by replacing contact frequencies S(Cij) by interface residue frequencies:

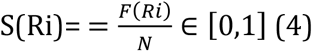

where F(Ri) denotes the frequency of residue i at the protein-protein interface (distance between heavy atoms lower than 5 Å) in the set of *N* decoys. The two related scores are respectively defined by:

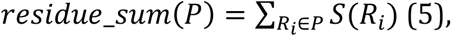

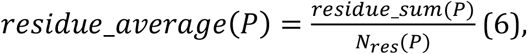

where S(Ri) is the relative frequency of interface residue Ri in the docking set of N decoys 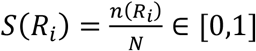 and *N_res_*(*P*) denotes the number of interface residues in pose *P*. Here, interface residues are simply those involved in contacts at the interface.

Note that is it possible to compute the contact and residue frequencies (equations 1 and 4) on a given set of docking poses and then to evaluate another set of docking poses (with equations 2, 3, 5 and 6).

### Clustering

We implemented the BSAS clustering procedure (Basic Sequential Algorithmic Scheme) ^32^ to reduce the structural redundancy of docking poses. The principle of BSAS is the following. Docking poses are ranked according to a score in decreasing order. The pose with the highest score initiates the first clusters. The other poses are sequentially compared to already clustered poses: they are included in a cluster if they are within a given cut-off of cluster members, otherwise they initiate a new cluster. At the end of the process, the pose with the highest score in each cluster is the representative of each cluster. In order to allow a fast clustering process, we do not compute the RMSD between ligand atoms. Instead we use a distance cut-off between the centers of mass of the ligands, here set to 8 Å.

### Evaluation

The top 2,000 solutions according to the ZDOCK native scoring function were rescored using the rescoring schemes detailed below. We monitored the presence of near-native docking hits (interface Cα RMSD < 2.5 Å) in the top 10 solutions after re-ranking. Each protein-protein complex with a near-native docking hit in the top 10 solutions is counted as a success.

### Implementation

The consensus-based rescoring functions are implemented in python code accessible on GitHub. All the scripts necessary to reproduce the results shown in this article are available at: https://github.com/MMSB-MOBI/CHOKO.

## RESULTS

### Quality of Decoys

In the initial data set of 176 protein-protein complexes, ZDOCK was able to generate at least one near-native docking hit (interface Cα RMSD < 2.5 Å) in the first top 2,000 solutions for 90 protein-protein complexes. These 90 protein-protein complexes thus constitute our reference data set for the rest of the study. We explore if and how consensus-based rescoring is efficient at scoring the decoys of these 90 protein-protein complexes.

### Evaluation of Different Consensus-Based Rescoring Functions

First, we compare the four versions of consensus-based rescoring functions: either contact-based or residue-based, with or without interface size normalization. Here, we remind the reader that the score based on contact frequencies with size normalization is the scheme proposed in CONSRANK ^26^. We estimate the performance by counting the number of successes, i.e., number of complexes with at least one near-native hit (interface Cα RMSD < 2.5 Å) in the first 10 solutions after rescoring. We also tested the effect of varying the subset of docking poses used to compute the contact and residue frequencies (equations 1 and 4): we used either the first 50, 100, 1,000 or 2,000 first poses provided by ZDOCK, referred as the frequency set. In any case, we rescored the first 2,000 poses provided by ZDOCK.

The results of this evaluation are shown in Figure 1. We can see that rescoring functions that take into account the interface size (Contact_Sum and Residue_Sum) constantly outperform the rescoring functions that disregard the interface size (CONSRANK, Residue_Average). The size of the subset used to compute contact and residue frequencies (equations 1 and 4) has a major influence on the number of successes. Indeed, ZDOCK solutions are ranked by the ZDOCK native scoring function; hence the top of the list is, in many cases, enriched in near-native docking hits. Estimating contact and residue scores on a reduced subset of poses at the top of the list is logically more efficient. On the contrary, estimating contact and residue scores from the full list leads to a loss of information, and worsens the prediction. In the best settings tested here, estimating the scores on the first 50 solutions to rescore the first 2,000 solutions allows to reach a number of successes equal to 27 with the Residue_Sum scoring function, *versus* 20 for the CONSRANK scheme. It is thus possible to rescore large sets of docking poses using consensus-based scoring functions, with better performance than the commonly used CONSRANK scheme.

**Figure 1.**
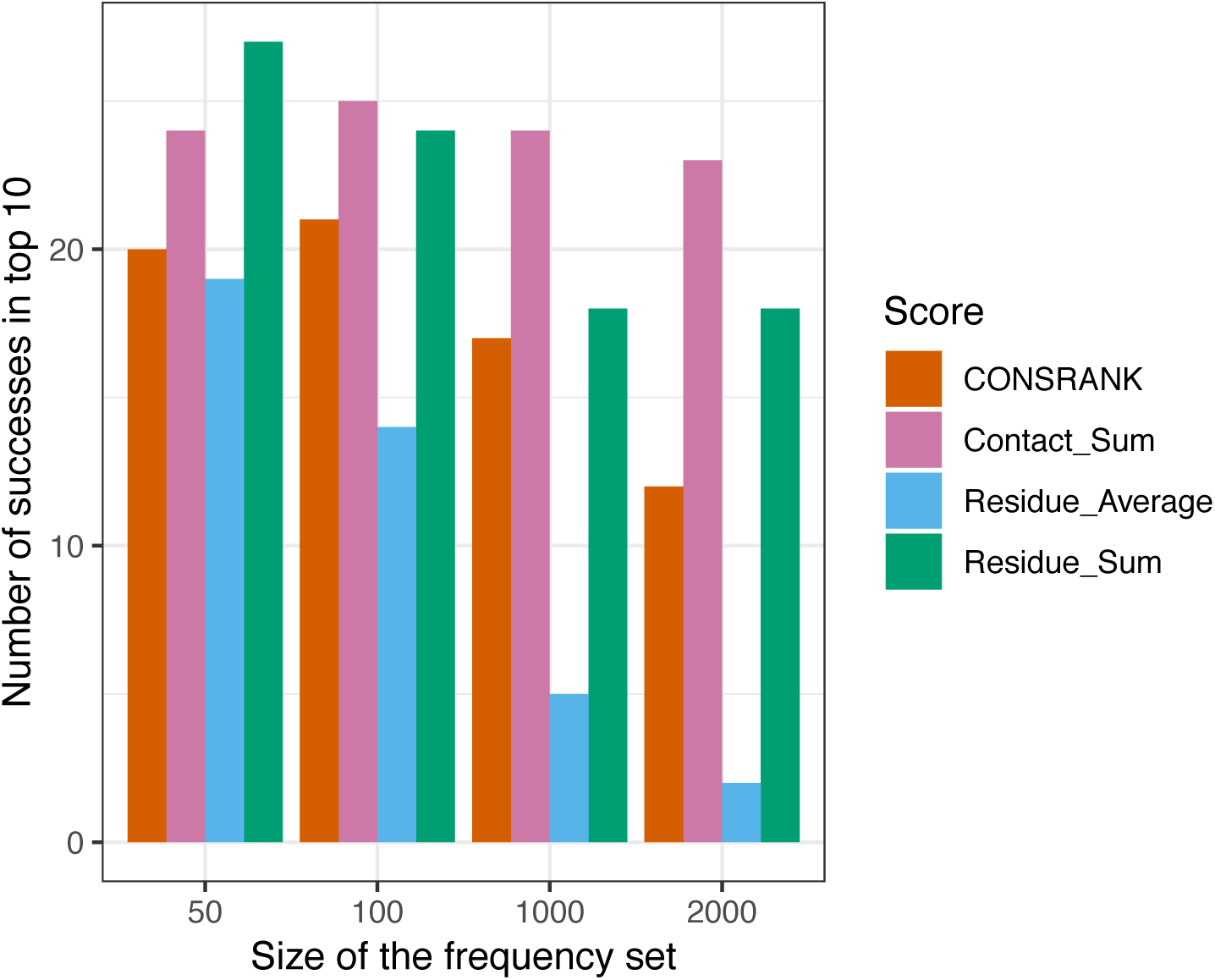
Number of successes after rescoring the first 2,000 solutions of ZDOCK. The size of the frequency set refers to the set of poses used to compute the residue and contact scores from equations 1 and 4.

### Combination With ZDOCK Native Scoring Function

In this section, we explore how to combine rescoring functions with the native scoring function of ZDOCK. Out of the 90 protein-protein complexes with at least one near-native docking hit in the top 2,000 solutions, the ZDOCK native scoring function identifies 29 successes, i.e., 29 complexes with at least one near-native docking hit in the top 10. This is indeed better than the four consensus-based rescoring functions tested here. One could wonder if it is then possible to improve the initial prediction of ZDOCK using rescoring.

We tested a combination of ZDOCK poses and rescored poses by combining the first N1 poses of ZDOCK with the first N2 poses after rescoring, with N1+N2=10, and no redundancy. Again, we vary the subset of docking poses used to compute the contact and residue frequencies with equations 1 and 4 (frequency set = top 50, 100, 1,000 or 2,000 poses) and in any case, we rescore the first 2,000 solutions provided by ZDOCK using equations 2, 3, 5 and 6. We estimate the performance by counting the number of successes, i.e., number of complexes with at least one near-native docking hit (interface Cα RMSD < 2.5 Å) in the first 10 solutions.

The results of this evaluation are shown in Figure 2. Regardless of the size of the frequency set, the best combination is always obtained with the Residue_Sum scoring function. Combining the first 6 ZDOCK poses with the first 4 Residue_Sum rescored poses, and estimating the frequencies on the full set of 2,000 poses (bottom right panel in Figure 2) allows to reach a number of successes equal to 32, which is more than Residue_Sum alone (18 successes) and better than ZDOCK alone (29 successes). It is thus possible to improve the native results of ZDOCK by a simple combination of poses. It is interesting to note that, in this situation, the information about residues is more efficient in rescoring than the information about pairwise contacts.

**Figure 2.**
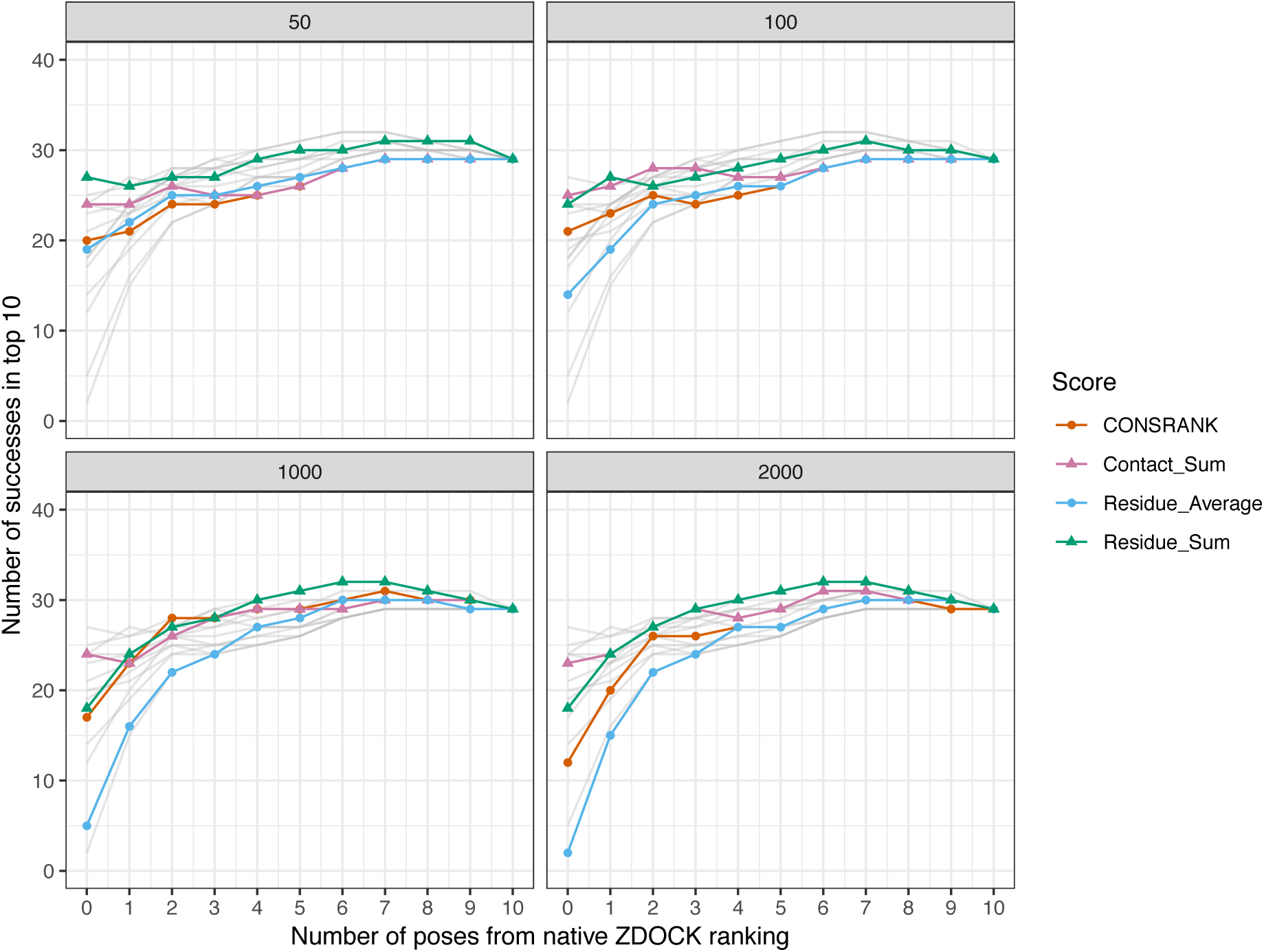
Number of successes after combination with the ZDOCK native scoring function. Each panel corresponds to different frequency sets, i.e., sets of poses used to compute the residue and contact scores from equations 1 and 4. In any cases, the first 2,000 solutions of ZDOCK are rescored. Grey lines represent data from other panels for comparison.

### Combining Clusters

Clustering is classically used to improve the performance, by reducing the structural redundancy of docking solutions ^33–35^. In this section, we explore how to combine rescoring functions with the native scoring function of ZDOCK, in conjunction with structural clustering. We used the BSAS clustering algorithm, which takes into account the different scores, since clusters are initiated according to the scores.

We have tested a combination of clusters. On the one hand, we computed clusters from the poses ranked by their initial ZDOCK scores. On the other hand, we computed clusters from poses reordered after consensus rescoring. We then combine the representative poses of the first N1 ZDOCK clusters, with the representative poses of the first N2 poses after rescoring, with N1+N2=10. We estimate the performance by counting the number of successes, i.e., number of complexes with at least one near-native hit (interface Cα RMSD < 2.5 Å) in the first 10 solutions.

The results of this evaluation are shown in Figure 3. The best combination is obtained using the Contact_Sum score, which constantly outperforms CONSRANK, regardless of the frequency set. When using the first 1,000 poses to estimate the frequencies (bottom left panel in Figure 3), the combination of five ZDOCK clusters and five Contact_Sum clusters achieves a number of successes equal to 38, which is better than ZDOCK alone (34 successes) and Contact_Sum alone (34 successes). Contrary to what was observed in simple pose combination (Figure 2), the most efficient rescoring scheme when dealing with clusters is based on contact frequencies, not residue frequencies. It seems that, after structural clustering, the information of pairwise contacts, which is more precise than residues, becomes more useful in discrimination.

**Figure 3.**
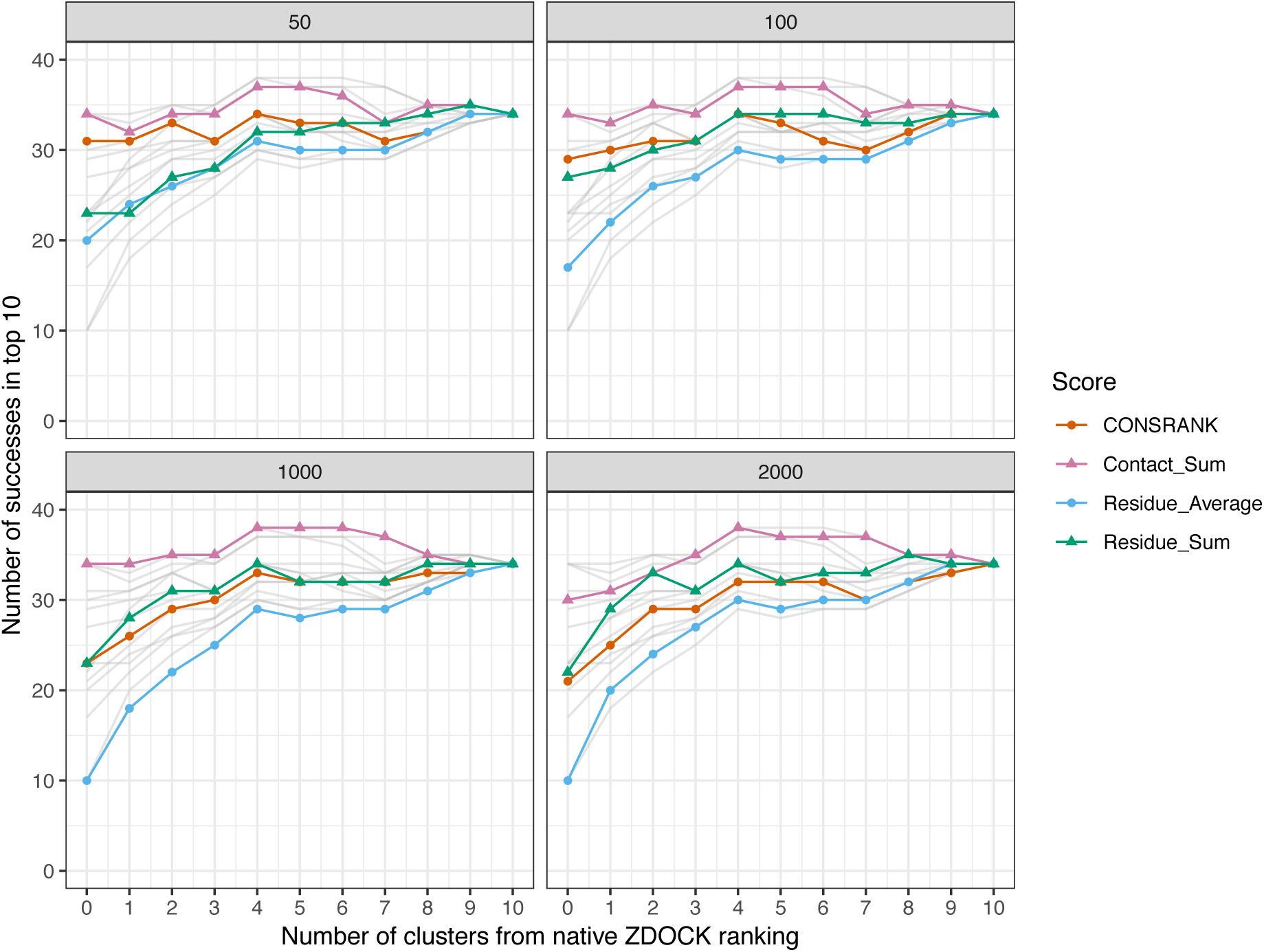
Number of successes after combination of clusters with the ZDOCK native scoring function. Each panel corresponds to different frequency sets, i.e., sets of poses used to compute the residue and contact scores from equations 1 and 4. In any cases, the first 2,000 solutions of ZDOCK are rescored.

### Illustrative Examples

To complete this study, in this section, we present examples to illustrate the asset of consensus rescoring when used in combination with the native ZDOCK scoring function. We used the results generated using the Contact_Sum function, with frequencies estimated on the first 1,000 poses, and combined five ZDOCK clusters and five consensus clusters. As explained in the previous section, this setting allows to reach 38 successes. We present four examples from the ZDOCK decoy set where the use of consensus rescoring is critical in Figure 4. For all these protein-protein complexes, no near-native docking hit is observed in the first ten clusters of ZDOCK (or in the first ten poses). The use of Contact_Sum rescoring in conjunction with clustering allows the identification of near-native docking in the top 10. In every case, these near-native poses do not belong to the top of the ZDOCK initial list: they are ranked 355 for 1AVX, 1568 for 1EAW, 250 for 1XQS and 606 for 1E6E. These examples highlight the usefulness of consensus-based rescoring to rescue poses with poor initial ranks.

**Figure 4.**
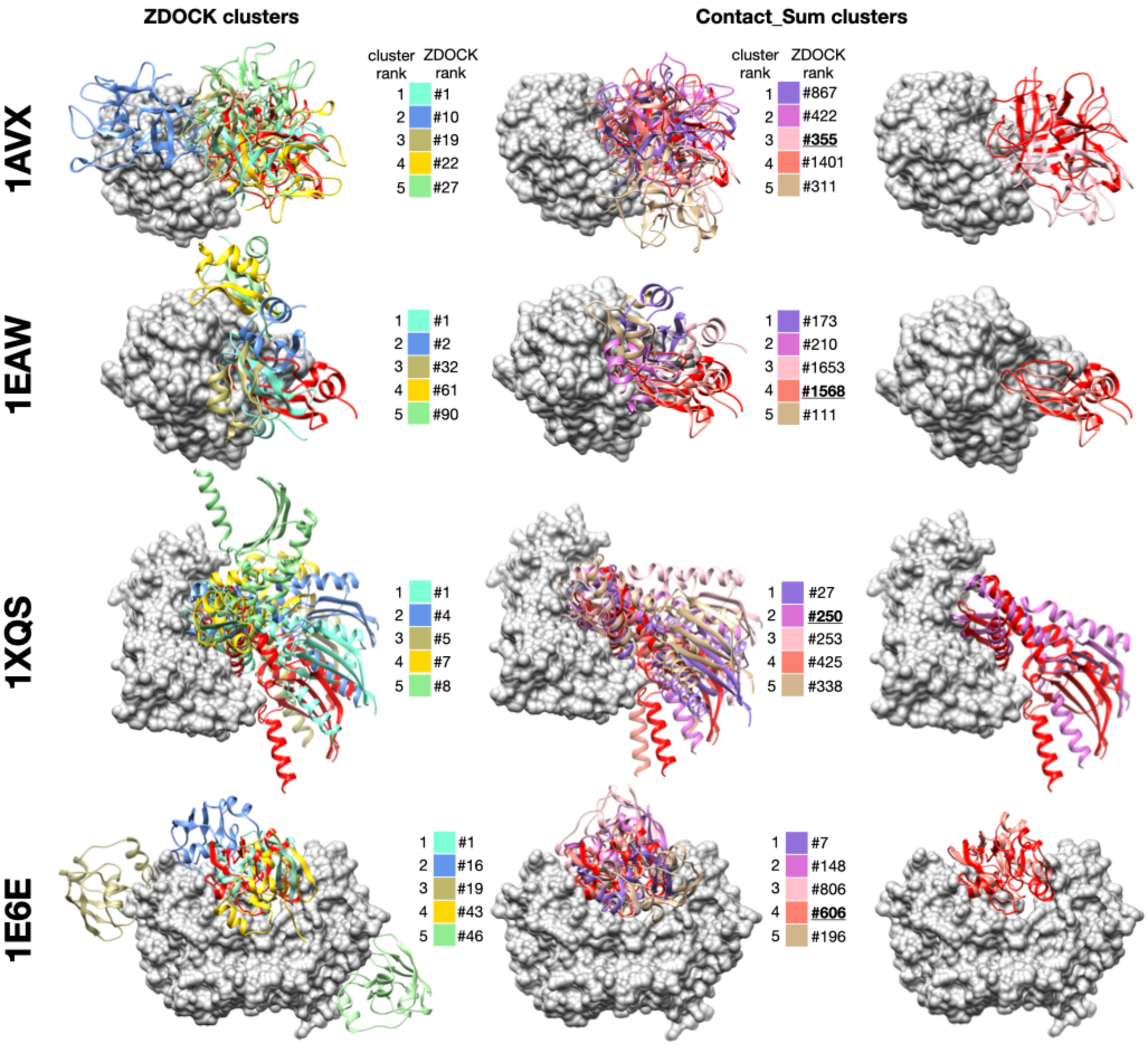
Examples of successful combination of ZDOCK clusters and consensus-based clusters. For each protein-protein complex, the receptor protein is represented as a gray surface and the ligand as a red ribbon. Left column: representative poses of the first five clusters generated with ZDOCK native scoring function, middle column: representative poses of the first five clusters generated with the Contact_Sum rescoring function, right column: superimposition between near-native docking hit and native structure. For each representative pose, initial ZDOCK rank is indicateed next to the color legend. Near-native poses are underlined. 1AVX ^36^:complex between the porcine trypsin (gray) and soybean inhibitor (red), 1EAW ^37^: complex between the catalytic domain of serine proteinase MT-SP1 (gray) and bovine inhibitor (red), 1XQS ^38^: complex between the human Hsp70 binding protein 1 (gray) and Hsp70 (red), 1E6E ^39^: complex between NADPH:adrenodoxin oxidoreductase (gray) and adrenoxin (red).

## CONCLUSION

We have implemented four variants of consensus-based rescoring functions, including the CONSRANK method, and tested them on the rescoring of large sets of docking poses of the ZDOCK benchmark. In this context, consensus scores that do take into account the size of the interfaces are in general more efficient than those that normalize by the size interface. The initial performances of ZDOCK are improved by combining the native scoring function with consensus-based rescoring.

## ACKNOWLEDGEMENTS

This work was financially supported by the “PHC Sakura” program, implemented by the French Ministry of Foreign Affairs, the French Ministry of Higher Education and Research and the Japan Society for Promotion of Science.

This project has received funding from the European Union’s Horizon 2020 Framework Programme for Research and Innovation under the Specific Grant Agreement No. 785907 (Human Brain Project SGA2).

